# Activation of Kiss1 neurons in the preoptic hypothalamus stimulates testosterone synthesis in adult male mice

**DOI:** 10.1101/2020.12.03.410878

**Authors:** Elisenda Sanz, Jonathan C. Bean, Richard D. Palmiter, Albert Quintana, G. Stanley McKnight

## Abstract

Kisspeptin-expressing neurons in the rostral periventricular region of the third ventricle (RP3V) play an essential role in female reproduction. However, adult male mice were reported to have very few Kisspeptin-expressing neurons in the RP3V compared to females. This led to the hypothesis that Kiss1^RP3V^ neurons are responsible for the ability of females, but not males, to generate a surge of LH, triggering ovulation and steroid synthesis in the female. Using mouse genetics and cell type-specific gene expression analysis, we show that male mice harbor almost as many Kiss1^RP3V^ neurons as the female and that gene expression in these neurons is very similar. Specific activation of male Kiss1^RP3V^ neurons expressing viral-encoded hM3Dq caused a surge in serum testosterone levels. These results demonstrate that Kiss1^RP3V^ neurons are present in the adult male and fully capable of regulating the hypothalamic/pituitary/gonadal axis. We suggest that these neurons may continue to play a role in reproductive behavior in adult male mice.

## INTRODUCTION

The brain contains several sexually dimorphic nuclei which are thought to coordinate differences in physiology and behavior between males and females. Within the rodent hypothalamus, the anteroventral periventricular nucleus (AVPV), a constituent of the rostral periventricular region of the third ventricle (RP3V), is thought to display sexually dimorphic neuronal populations including tyrosine hydroxylase (*Th*)-, or kisspeptin (*Kiss1*)-expressing cells (Clarkson & Herbison, 2006; Simerly et al., 1997). Kiss1 neurons are critical to the reproductive system. Humans and mice deficient in different components of the kisspeptin signaling system present with hypogonadotropic hypogonadism, infertility and absence of puberty (D’anglemont De Tassigny et al., 2007; De Roux et al., 2003; Funes et al., 2003; Seminara et al., 2003). Two major Kiss1 neuronal populations have been reported in the mouse hypothalamus. One population is located in the RP3V and the other in the arcuate hypothalamus (ARC) (Gottsch et al., 2004), and they both project to GnRH neurons and regulate the hypothalamic-pituitary-gonadal (HPG) axis. *Kiss1* mRNA expression in the RP3V, but not in the ARC Kiss1 (Kiss1^ARC^) neurons, differs between the sexes with the male RP3V appearing to harbor a greatly reduced number of Kiss1-expressing cells when compared to females as assayed by standard *in situ* hybridization techniques (Kauffman, 2010). In adult female mice the Kiss1^RP3V^ neurons are thought to stimulate the preovulatory surge of GnRH and LH in response to estrogen (Clarkson & Herbison, 2016). Adult male mice do not respond to either estrogen or testosterone with an LH surge and this has been attributed to the loss of Kiss1 neurons. The physiological role for male Kiss1^RP3V^ neurons has been hypothesized to involve the triggering of the neonatal surge of testosterone after which this population of male Kiss1 neurons was hypothesized to undergo apoptosis (Clarkson & Herbison, 2016). In rodents, administration of either androgen or estrogen to newborn females causes a male-like pattern of reduced *Kiss1* neurons in the RP3V and an absence of cyclic LH surges in adult females. In males, neonatal castration leads to feminization of Kiss1^RP3V^ neurons and the mice retain the ability to generate estrogen-induced surges as adults (Homma et al., 2009). Several mechanisms including postnatal programed cell death and epigenetic silencing of the *Kiss1* gene have been proposed to underlie this steroid-mediated sex-specific difference (Semaan & Kauffman, 2013). We have reexamined the male Kiss1^RP3V^ neuronal population in adults using more sensitive techniques and established that these Kiss1 neurons are still present in the male RP3V and continue to express Kiss1 as well as other marker genes that identify them as similar to their female counterparts. Furthermore, we provide evidence that these neurons remain functionally connected within the HPG axis and are capable of initiating a rapid increase in testosterone production.

## RESULTS

### Adult male and female Kiss1^RP3V^ neurons are similar in number

Sex-dependent differences in Kiss1^RP3V^ neurons arise at postnatal day 12 and lead to an evident sexual dimorphism in the adult when assessed by *in situ* hybridization for *Kiss1* mRNA (Semaan & Kauffman, 2013). To explore whether this difference is due to a downregulation of *Kiss1* mRNA in *Kiss1*-expressing cells or to the cell death of this neuronal population in male mice, we crossed mice expressing Cre recombinase under the *Kiss1* promoter (Kiss1-Cre mice) (Gottsch et al., 2011) to mice expressing the fluorescent protein TdTomato in a Cre-dependent manner (Rosa26-flox-stop-TdT mice) (Madisen et al., 2010) (Figure 1A). The expression of TdTomato in the RP3V of male and female Kiss1-Cre:Rosa26-flox-stop-TdT mice is shown (Figure 1B). Imaging of sections containing the RP3V in both male and female mice showed a prominent number of TdTomato-expressing cells, suggesting that the apparent reduction in the number of *Kiss1*-expressing cells in the RP3V of adult male mice is not due to cell death. Cell counts for RP3V TdTomato positive neurons in the male were only slightly reduced compared to those in the female (Figure 1C).

**Figure 1.**
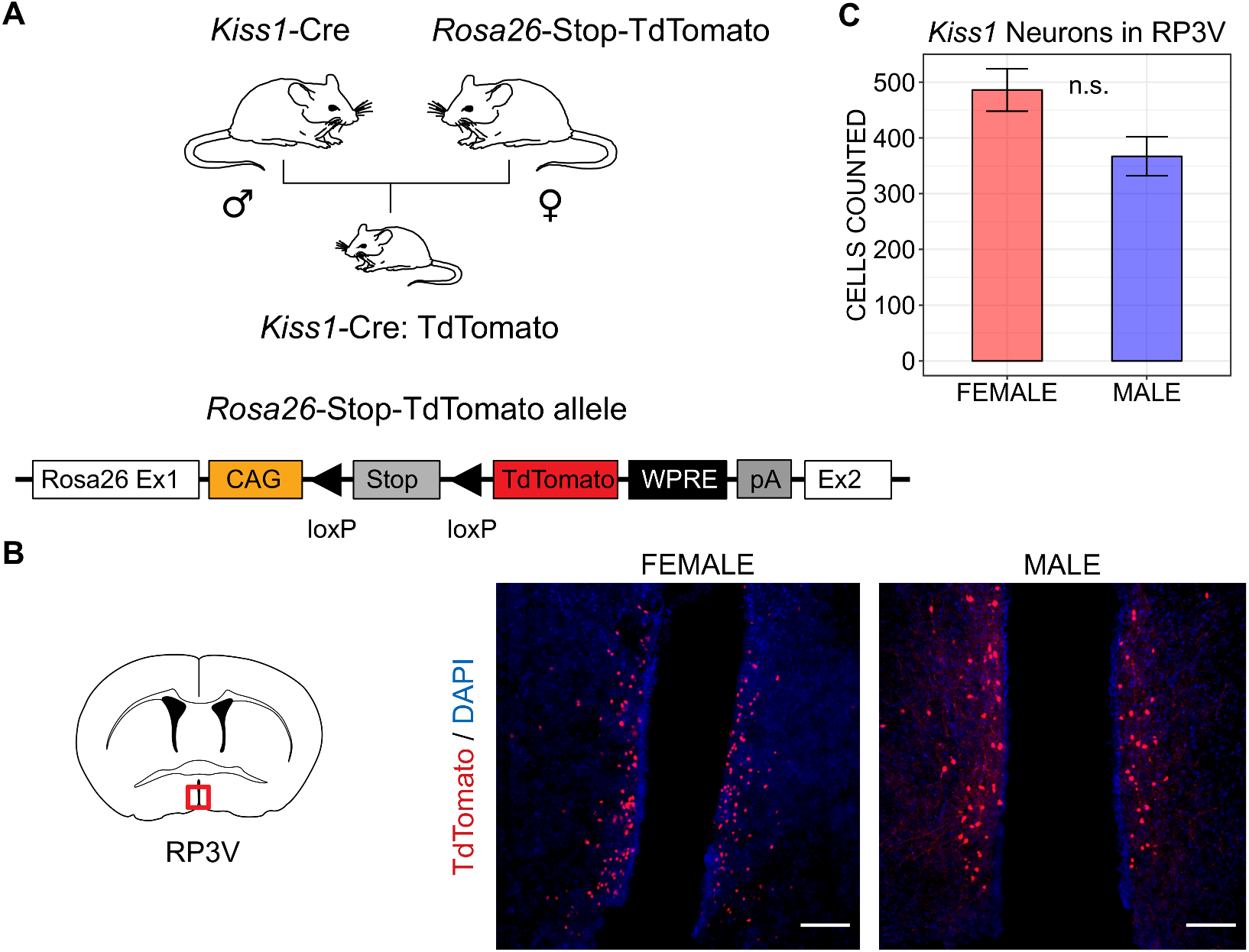
Adult male and female mice have similar numbers of Kiss1^RP3V^ neurons. (A) Kiss1-Cre:TdTomato mice were the progeny of a male Kiss1-Cre and the Ai14 TdTomato reporter strain available from Jackson Labs. The TdTomato allele is expressed after Cre recombinase removal of the Stop signals as depicted in the schematic. (B) Cartoon of a coronal brain section with the location of the RP3V outlined by a red box. TdTomato fluorescence of a representative coronal section containing the RP3V in female and male adult mice. DAPI staining of the nuclei is also shown. Scale bars: 100 μm. (C) Sections (30 μm) through the RP3V of 2 male and 2 female Kiss1-Cre:TdTomato mice were obtained and TdTomato positive cells were counted in every 4^th^ section. The total number of cells counted is shown for females and males.

### RiboTag analysis of marker transcripts in the Kiss1^ARC^ and Kiss1^RP3V^ neurons in male and female hypothalamus

We compared the actively translated mRNAs in male and female Kiss1^RP3V^ neurons using a ribosome-tagging (RiboTag) strategy developed in our lab (Sanz et al., 2019; Sanz et al., 2009). To make this comparison, we crossed the Kiss1-Cre mice to RiboTag mice that will express an HA-tagged 60S ribosomal protein (Rpl22-HA) in response to Cre recombination. The wild type untagged exon 4 is removed from the gene by Cre-dependent recombination allowing the HA-tagged exon 4 to be translated as depicted in Figure 2A. Both the RP3V and ARC regions of the hypothalamus were isolated by dissection and the HA-tagged polyribosomes were immunoprecipitated (IP) using anti-HA monoclonal antibody and protein A/G magnetic beads as described (Sanz et al., 2019). Total RNA samples prepared from both the input homogenates and the IP-isolated polyribosomes were then analyzed by RNA-Seq (Figure 2B).

**Figure 2.**
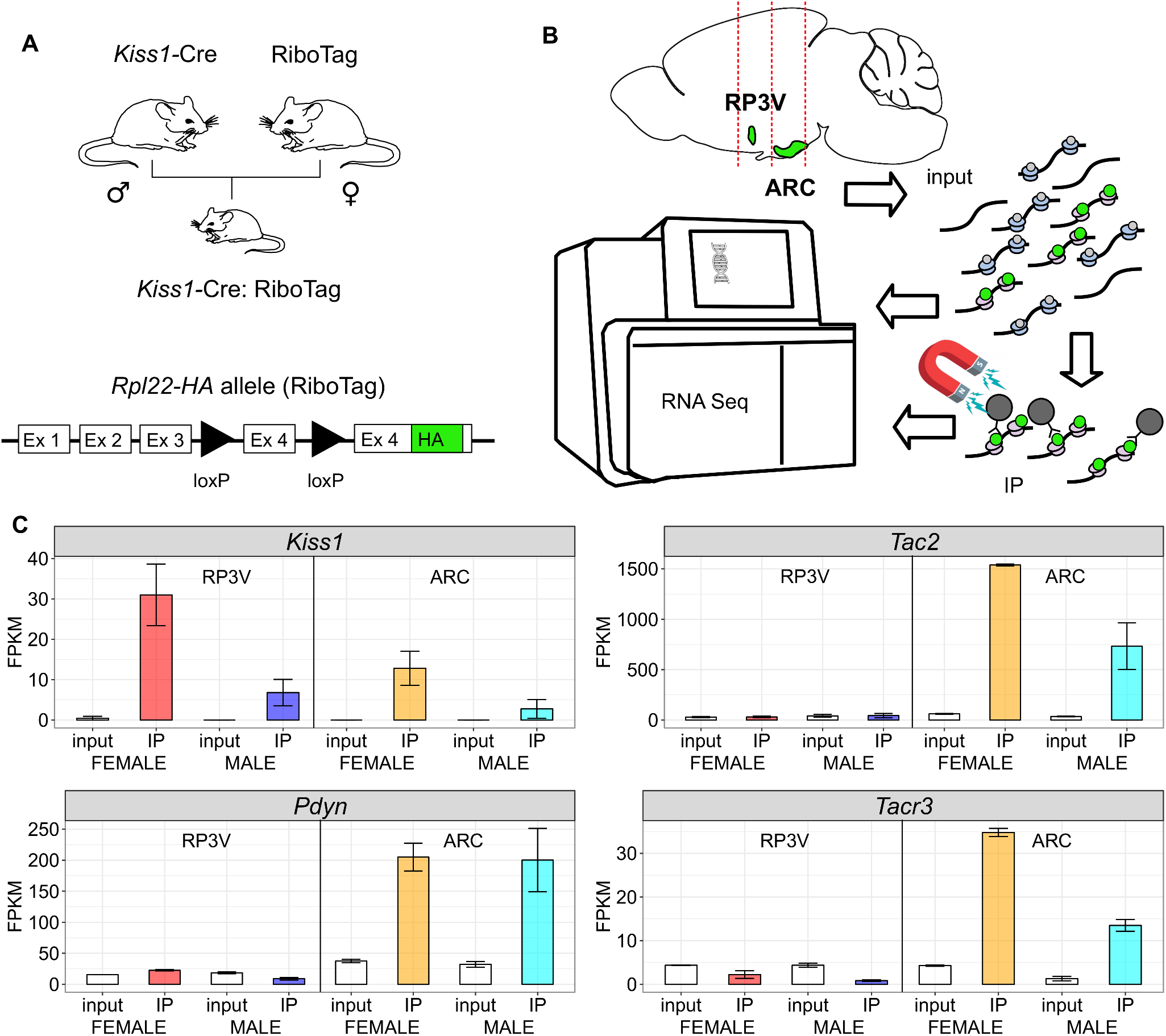
Expression of marker genes in Kiss1^RP3V^ and Kiss1^ARC^ neurons in male and female adult mice. (A) Kiss1-Cre:RiboTag mice were the progeny of male Kiss1-Cre and female RiboTag mice. The Rpl22-HA allele contains a loxP-flanked wild type exon 4 which is replaced by an HA-tagged exon 4 after Cre recombination. (B) Mouse brains were sliced into 2mm coronal slices that anatomically separated the ARC and RP3V regions of the hypothalamus. The 2 mm punches of ARC and RP3V tissue were homogenized, used to immunoprecipitate (IP) HA-tagged polysomes, and then RNA was extracted for RNA-Seq. A sample of the total homogenate was also used to prepare an input RNA sample for RNA-Seq for comparison to the IP RNA. Punches from four animals were pooled for each RiboTag isolation and two separate groups of mice were analyzed. (C) Expression of Kiss1 and the Kiss1^ARC^ neuronal markers, *Tac2, Pdyn*, and *Tacr3* are shown comparing male and female RP3V and ARC. Expression is given as FPKM (Fragments Per Kilobase per Million reads) and the range of the 2 groups of 4 animals each is shown by error bars.

The Kiss1^ARC^ neurons from both male and female mice are enriched (IP greater than input) for transcripts from the *Kiss1, Tac2*, *Pdyn* (dynorphin) and *Tacr3* genes as expected. The levels of *Kiss1, Tac2*, and *Tacr3* mRNAs were substantially lower in the male ARC but both sexes expressed similar levels of *Pdyn* as shown in Figure 2C. The Kiss1^RP3V^ neurons from both males and females were enriched in *Kiss1* mRNA (Figure 2C) but the males expressed 4fold lower levels which may explain why *in situ* hybridization failed to detect many of the male Kiss1^RP3V^ neurons. As expected, marker genes for astrocytes, oligodendrocyctes, and microglia were depleted in IP samples whereas housekeeping genes such as *Actb* (β–actin) and *Gapdh* were expressed at similar levels in both input and IP as shown (Figure 2–figure supplement 1). The female Kiss1^ARC^ neurons have been shown to be predominantly glutamatergic (Nestor et al., 2016) whereas up to 70% of female Kiss1^RP3V^ neurons are GABAergic (Cravo et al., 2011; Nestor et al., 2016). Expression of the GABA synthetic enzyme, *Gad2*, and the glutamate transporter, Vglut2 (*Slc17a6*), are consistent with these published results for the female and show that the male Kiss1^RP3V^ neurons share similar expression patterns (Figure 2—figure supplement 1).

### Male and female Kiss1^RP3V^ neurons preferentially express a similar signature of neuropeptide, receptor, and other marker genes

We next examined whether the same neuropeptide and receptor genes that are enriched in female Kiss1^RP3V^ neurons are also expressed and enriched in male Kiss1^RP3V^ neurons. Previous studies have reported that galanin (*Gal*)(Porteous et al., 2011), proenkephalin (*Penk*)(Porteous et al., 2011), tyrosine hydroxylase (*Th*)(Clarkson & Herbison, 2011), estrogen receptor (*Esr1*)(*Cravo* et al., 2011), progesterone receptor (*Pgr*) (Stephens et al., 2015) and the ghrelin receptor (*Ghsr*)(Frazao et al., 2014) mRNAs are all co-expressed with *Kiss1* mRNA in at least a subset of neurons in the female RP3V. Our results demonstrate that these mRNAs are all enriched (IP/Input ≥2) in both female and male Kiss1^RP3V^ neurons (Figure 3). RNA-Seq analysis also identified the enriched expression of other neuropeptide and receptor mRNAs in both male and female Kiss1^RP3V^ neurons including neuromedin U (*Nmu*) and neurotensin (*Nts*) (Figure 3). Expression of the androgen receptor (*Ar*) was slightly enriched (1.3 in females and 1.7 in males) compared with the input but above the background of the RiboTag assay. These results indicate that the male Kiss1^RP3V^ neurons are quite similar to their female counterparts.

**Figure 3.**
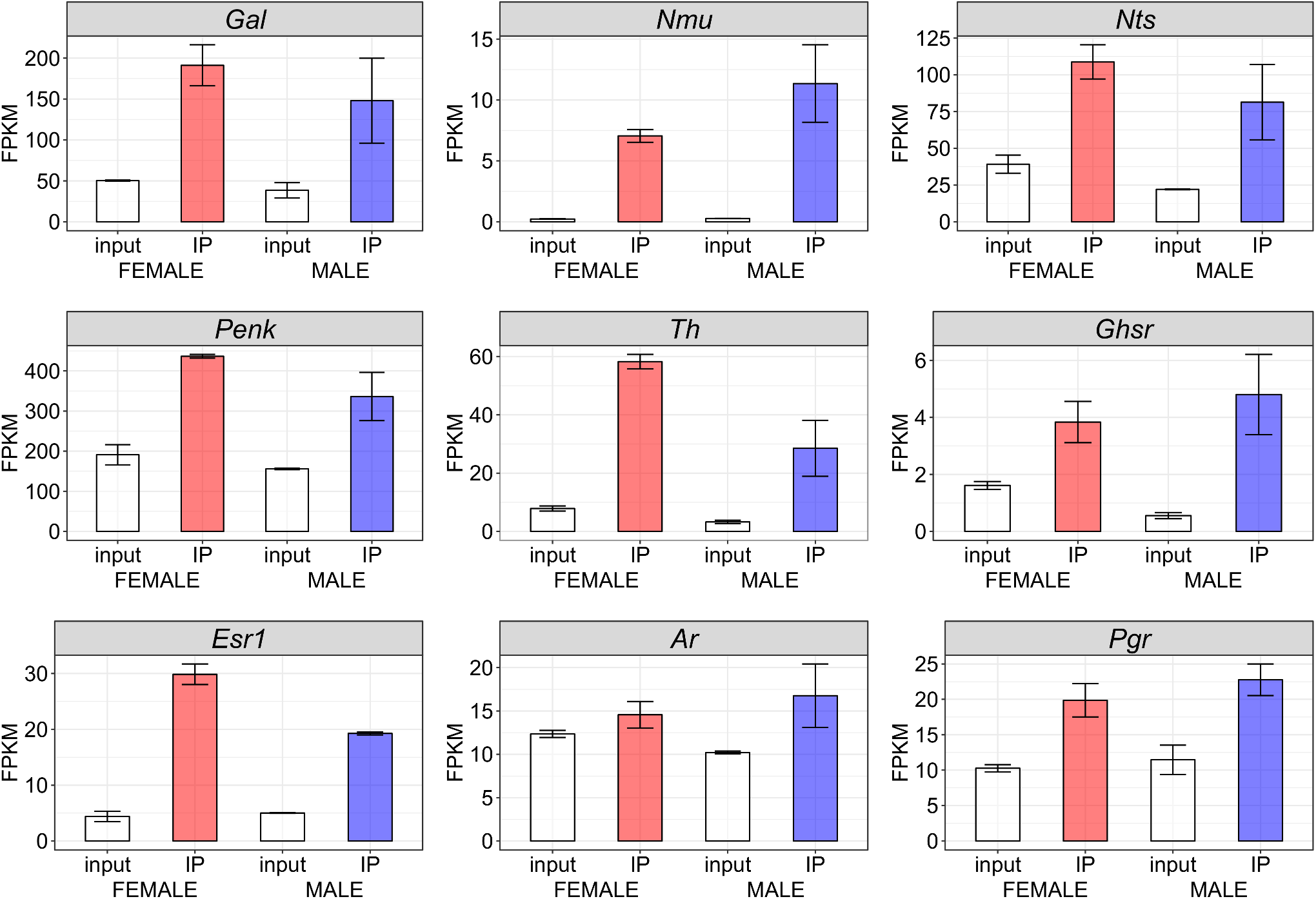
Male and female Kiss1^RP3V^ neurons express and translate a similar signature of transcripts. The RNA-Seq results show enriched expression from the neuropeptide genes, *Gal*, *Nmu, Nts*, and *Penk*. The tyrosine hydroxylase gene (*Th*), the ghrelin receptor gene (*Ghsr*) and the steroid receptor genes (*Esr1, Ar*, and *Pgr*) are all expressed in the Kiss1^RP3V^ neurons although *Ar* is only modestly enriched. Genes that are not expressed in the HA-tagged neurons will still show up in the IP samples as background with an IP/Input ratio ranging from 0.1 to 0.4 as shown for glial marker genes in Figure 2-Supplement 1. Expression levels are normalized to FPKM and the range of the 2 samples (4 mice pooled in each sample) is shown by the error bars.

By crossing Kiss1-Cre mice with RiboTag mice we enabled expression of RiboTag in adult neurons that might have expressed Kiss1-Cre during development but are no longer expressing Cre recombinase in the adult. To confirm that the marker genes we identified are still expressed in adult Kiss1^RP3V^ neurons we repeated the RiboTag analysis by injecting the Cre-dependent RiboTag (AAV-DIO-Rpl22HA) as described (Sanz et al., 2015). The results shown in Figure 3–figure supplement 1 confirm that the neuropeptides, neuromodulators, and receptors that we identified in our comparison of male and female Kiss1^RP3V^ neurons using the RiboTag mice continue to be expressed in the adult Kiss1 neurons.

Neuronal subtypes can be distinguished by the specific neuropeptides, receptors, and transcription factors that they express and this approach has been used to compare and group neurons after single cell RNA-Seq. We used a curated list of neuropeptides and receptors (Moffitt et al., 2018) and an atlas of mouse transcription factors (Zhou et al., 2017) to extend our analysis and make an unbiased comparison between male and female Kiss1^RP3V^ neurons. For each group of genes, we filtered for those expressed at 5 FPKM or more in either the input or IP of male or female samples and then calculated the enrichment (IP/Input) for the neuropeptides and receptors that passed this filter. We also examined the quantitative expression of each gene using the IP values for genes that were enriched (IP/Input) ≥ 1. The graphs demonstrate that the signature of neuropeptide genes expressed in Kiss1^RP3V^ neurons is similar in terms of both enrichment and quantitative expression (Figure 3–figure supplement 2, Panels A,B). Analysis of receptor mRNAs (Figure 3-figure supplement 2, Panels C,D) and transcription factor mRNAs (Figure 3-figure supplement 2, Panel E) also illustrate the similarity of male and female Kiss1^RP3V^ neurons. For the transcription factors we used an enrichment ≥1.5 as the filter.

### Response of male Kiss1 neurons to steroid hormone deprivation

Female Kiss1 neurons in both the ARC and RP3V express estrogen receptors and respond to estrogen with changes in mRNA expression and electrophysiological responses (Qiu et al., 2018; Zhang et al., 2015; Zhang et al., 2013). Estrogen inhibits the expression of *Kiss1* mRNA in ARC neurons as part of a feedback-inhibitory pathway. However, estrogen induces *Kiss1* mRNA and increases electrical excitability in Kiss1^RP3V^ neurons that synapse on GnRH neuronal cell bodies. The increase in *Kiss1* mRNA and activation of Kiss1^RP3V^ neurons is thought to stimulate the release of GnRH, triggering the surge of LH leading to ovulation (Clarkson & Herbison, 2016). The expression of *Kiss1* mRNA in the male has also been studied and castration induced Kiss1 mRNA in the Kiss1^ARC^ neurons and also inhibited Kiss1 mRNA expression in Kiss1^RP3V^ neurons (Smith et al., 2005). We used Kiss1-Cre:RiboTag mice to examine the global transcriptional phenotype of Kiss1 neurons in both the ARC and RP3V of intact and castrated males. Castration caused a dramatic stimulation of *Kiss1, Tac2*, *Pdyn*, and *Tacr3* mRNAs in the Kiss1^ARC^ neurons (Figure 4A). In the same intact and castrated mice we examined the ribosome bound mRNA transcripts in Kiss1^RP3V^ neurons.

**Figure 4.**
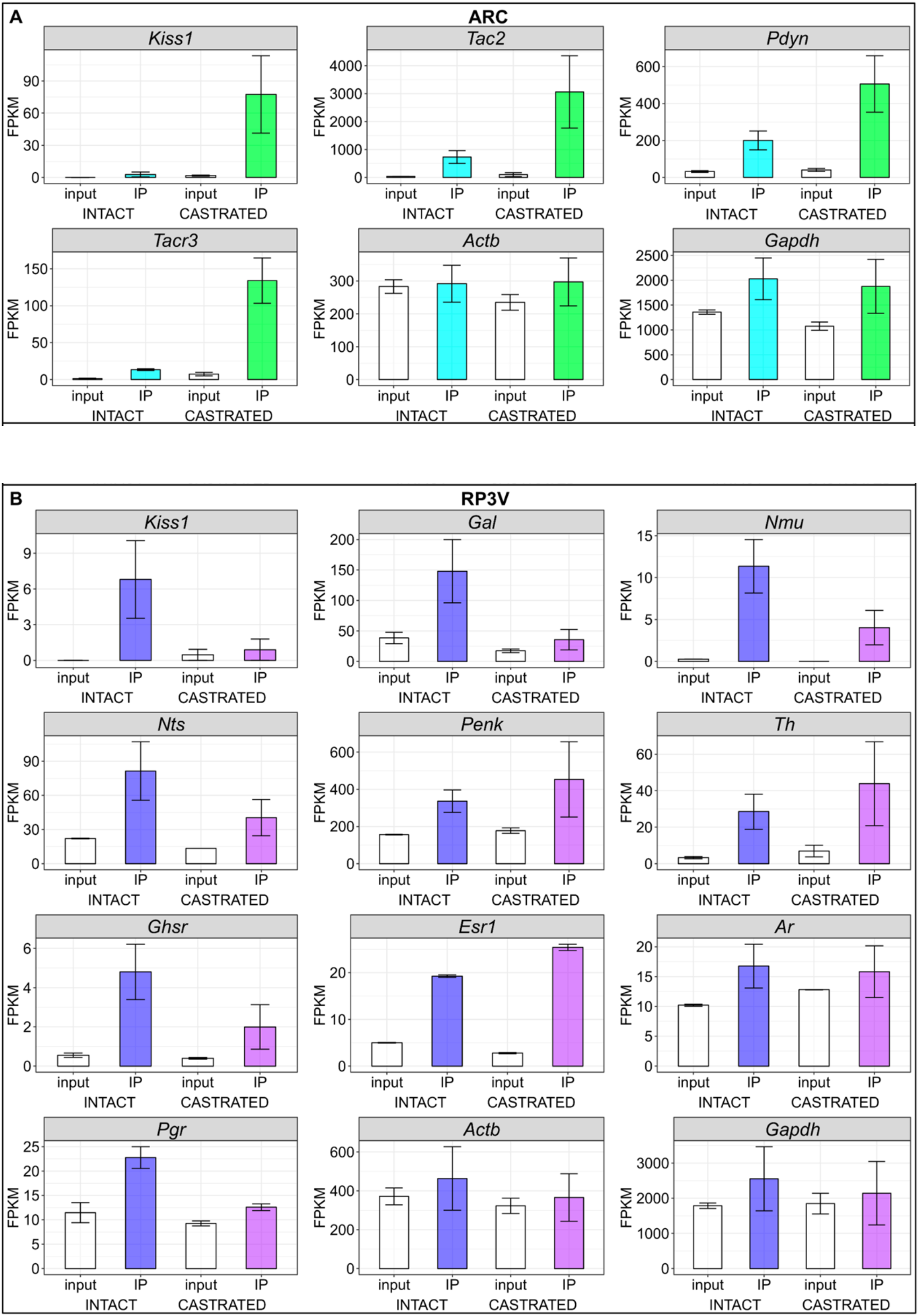
Androgen depletion by castration alters expression of Kiss1 and a subset of enriched marker genes in male Kiss1 neurons. Male Kiss1-Cre:RiboTag mice were castrated as adults and 4 weeks later were analyzed for mRNA expression by RiboTag and the results compared with male Kiss1-Cre:RiboTag controls (Intact). (A) Expression levels of *Kiss1, Tac2*, *Pdyn*, and *Tacr3* in the Kiss1^ARC^ neurons of intact and castrated mice are shown. For comparison the constitutively expressed gene transcripts for *Actb* and *Gapdh* are also shown. (B) Expression of *Kiss1* and other marker genes in Kiss1^RP3V^ neurons is shown for intact and castrated male mice. The constitutively expressed genes, *Actb* and *Gapdh* are shown for comparison. In both panels, the error bars show the range of values for 2 separate pools of mice, each containing tissues harvested from 4 animals. Values for intact males are the same as those shown in Figure 2.

Testosterone deprivation by castration decreased the expression of *Kiss1, Nmu, Nts, Gal, Pgr*, and *Ghsr* by 2-5 fold in Kiss1^RP3V^ neurons without affecting the expression of *Penk, Th, Esr1* or *Ar* mRNAs or the expression of housekeeping mRNAs, *Actb* or *Gapdh* (Figure 4B).

### Male Kiss1^RP3V^ neurons are capable of stimulating the HPG axis and producing a surge of circulating testosterone

To explore the functional role of the male Kiss1^RP3V^ neuronal population in the regulation of the HPG axis, we introduced a virus that expresses a Cre recombinase dependent activating DREADD, hM3Dq, fused to the fluorescent protein mCherry (AAV1-DIO-hM3Dq:mCherry) (Krashes et al., 2011). The AAV1-DIO-hM3Dq:mCherry virus was injected bilaterally into the RP3V of male *Kiss1-Cre* mice. (Figure 5A). Brain sections containing the RP3V and the ARC showed that the mCherry signal was restricted to the RP3V and that no expression of the activating DREADD was detected in the ARC (Figure 5B). Administration of the synthetic DREADD ligand, clozapine-N-oxide (CNO) to these mice resulted in a dramatic increase in serum testosterone within 1 hour whereas administration of CNO to control mice showed no increase in serum testosterone levels (Figure 5C, D). To confirm specific neuronal activation after CNO administration, *Kiss1-Cre* mice were unilaterally injected with AAV1-DIO-hM3Dq:mCherry into the RP3V and Fos staining was assessed 1 hour after CNO injection. RP3V sections showed a significant number of Fos-positive nuclei only on the side injected with the activating DREADD virus, confirming the specific activation of hM3Dq:mCherry-expressing neurons after CNO administration (Figure 5E).

**Figure 5.**
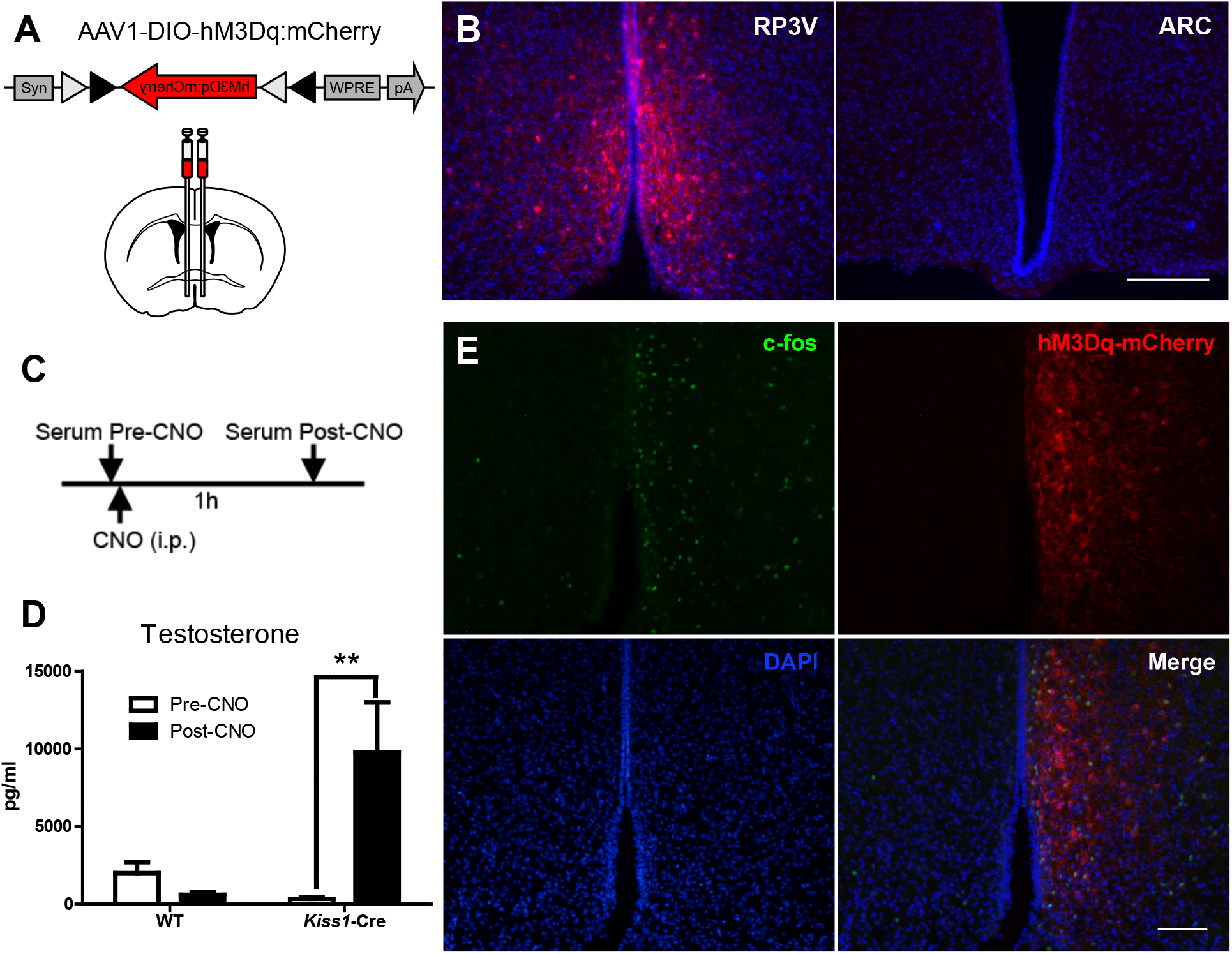
Activation of Kiss1^RP3V^ neurons using hM3Dq and CNO leads to a rapid surge in plasma testosterone. **(A)** A schematic of the AAV1-DIO-hM3Dq:mCherry virus is shown. The virus was injected bilaterally into the RP3V region of Kiss1-Cre mice. **(B)** Four weeks after injection the expression of mCherry is prominent in RP3V but not in the ARC of the same animals. Scale bar: 100 μm. (**C)** Diagram showing the time points for serum collection before and after i.p. administration of CNO. **(D)** Serum testosterone levels in WT and Kiss1-Cre mice (n=4) injected with AAV1-DIO-hM3Dq:mCherry pre and post injection with 1 mg/kg CNO. Data are presented as the mean ± SEM. Statistical analysis was performed using two-way ANOVA followed by Bonferroni posttest (**p<0.01). **(E)** Male Kiss1-Cre mice were injected unilaterally with AAV1-DIO-hM3Dq:mCherry and 4 weeks later CNO (1mg/kg) was administered. After 1 h, Fos was visualized by immunohistochemistry and mCherry was visualized by fluorescence allowing direct visualization of hM3Dq expression in the RP3V region of the hypothalamus. DAPI staining of DNA is shown to identify cell nuclei.

## DISCUSSION

The cellular and molecular players that control the HPG axis and regulate reproduction and behavior are incompletely understood. Hypothalamic Kiss1 neurons are thought to initiate puberty in both male and female rodents and regulate the ovulatory cycles in the female. Kiss1 neurons in the ARC are inhibited by estrogen and testosterone and this provides a negative feedback mechanism to regulate the HPG axis in both sexes. In contrast, Kiss1^RP3V^ neurons are stimulated by estrogen and have been hypothesized to provide a feed-forward signal to trigger GnRH neurons and initiate the ovulatory surge of LH (Oakley et al., 2009). Our understanding of the physiological role, if any, of Kiss1^RP3V^ neurons in the adult male rodent is limited and it has been suggested that these neurons undergo apoptosis after their early activation during the neonatal surge of testosterone that only occurs in the male (Clarkson & Herbison, 2016). Here, we used Kiss1-Cre mice to identify the adult population of Kiss1^RP3V^ neurons by activation of a TdTomato reporter and showed Kiss1^RP3V^ neurons were not lost during development in the male mouse as had been suggested based on *in situ* hybridization (Clarkson & Herbison, 2016). Combining this Kiss1-Cre with the RiboTag approach we identified actively translated mRNAs that are expressed and enriched in both RP3V as well as ARC Kiss1 neurons.

Analysis of neuropeptide, receptor, transcription factor, and neuromodulator expression in adult Kiss1^RP3V^ neurons demonstrates that both male and female Kiss1 neurons in this region are preferentially translating a similar collection of function-specifying mRNAs coding for neuropeptides, receptors, and transcription factors. However, *Kiss1* mRNA is expressed at a 4-fold lower level in male compared with female Kiss1^RP3V^ neurons providing a likely explanation for the inability of classical *in situ* hybridization techniques to accurately identify this population in the male. Kiss1^RP3V^ neurons from both sexes express enriched levels of the neuropeptide mRNAs: galanin (*Gal*), neuromedin U (*Nm*u), neurotensin (*Nts*), and enkephalin (*Penk*). They also selectively express the ghrelin receptor (*Ghsr*), estrogen receptor alpha (*Esr1*), and progesterone receptor (*Pgr*) mRNAs. The mRNA for the rate-limiting enzyme required for catecholamine synthesis, tyrosine hydroxylase (*Th*), is also significantly enriched in both male and female Kiss1^RP3V^ neurons in agreement with previous studies showing co-expression of *Th* and *Kiss1* in female RP3V neurons (Clarkson & Herbison, 2011). Our results are also in general agreement with a recent analysis of mRNA expression in the mouse preoptic area (POA) by single cell RNA-Seq, although this study did not compare male and female-specific expression for Kiss1^RP3V^ neurons (Moffitt et al., 2018).

In female rodents, removal of estrogen by ovariectomy increases expression of *Kiss1* mRNA in the ARC while inhibiting *Kiss1* mRNA expression in the RP3V but the mechanisms that elicit this bidirectional regulation are not completely understood. We compared the mRNA expression patterns of Kiss1 neurons from intact and castrated adult male mice and demonstrated that castration caused a marked stimulation of *Kiss1, Tac2, Pdyn*, and *Tacr3* mRNAs in the ARC as previously described (Smith et al., 2005) (Navarro et al., 2011). *Kiss1* mRNA expression in the male RP3V is also regulated by sex steroids and we find that castration caused a dramatic reduction in *Kiss1* mRNA in the RP3V. Castration also decreased the expression of a sub-group of Kiss1^RP3V^ neuron-enriched marker transcripts including *Gal, Nmu, Nts, Ghsr*, and *Pgr* without affecting expression of *Esr1, Ar*, or housekeeping genes (*Actb, Gapdh*).

Kiss1 neurons in the RP3V have been studied extensively in female rodents. The majority express the estrogen receptor, ERα (*Esr1*), and at least 30% of them project to the cell bodies of GnRH neurons in the POA where they provide an excitatory input that is thought to be responsible for the preovulatory surge of LH (Cravo et al., 2011; Kumar et al., 2015; Wang & Moenter, 2020). Despite their stimulatory function, 70% of the Kiss1^RP3V^ neurons appear to be GABAergic and only 20% are identified as glutamatergic (Cravo et al., 2011). Optogenetic activation of Kiss1^RP3V^ neurons in female mice stimulates the firing of GnRH neurons in brain slices and leads to a sustained increase in LH secretion *in vivo*. This Kiss1^RP3V^ neuron-induced surge of LH is completely dependent on kisspeptin expression (Piet et al., 2018). The LH surge in response to estrogen is also dependent on the expression of ERa in RP3V^Kiss1^ neurons which supports the hypothesis that Kiss1^RP3V^ neurons serve as the estrogen-dependent surge generator that triggers ovulation (Wang et al., 2019).

Our results demonstrate the existence of a population of Kiss1^RP3V^ neurons in the adult male that are very similar to the female Kiss1^RP3V^ neurons in terms of population size and enriched expression of marker neuropeptides, receptors and other marker genes. To begin to explore the physiological function of this population of Kiss1 neurons we determined whether the adult male Kiss1^RP3V^ neurons can stimulate the HPG axis. Expression of hM3Dq DREADD specifically in Kiss1^RP3V^ neurons and pharmacological activation with CNO evoked a dramatic elevation of circulating testosterone within 1 hour demonstrating that these Kiss1 neurons retain the ability to stimulate the HPG axis as do their female counterparts. Our results also suggest that the inability of estradiol to stimulate a surge of LH and testosterone in male mice cannot be explained by a loss of ERa or PR since Kiss1^RP3V^ neurons in the male express levels of these steroid receptor mRNAs similar to the female. The lack of an LH surge in adult male mice in response to estradiol also seems unlikely to be solely due to the decreased expression of kisspeptin since direct stimulation of male Kiss1^RP3V^ neurons is capable of stimulating the HPG axis. In female rodents there is strong evidence that inputs from the suprachiasmatic nucleus (SCN) gate the ability of estradiol to elicit an LH surge (Williams et al., 2011). Poling et al. (Poling et al., 2017) implanted mice with silastic capsules containing estradiol and showed that female Kiss1^RP3V^ neurons become active 2 days later just before lights out based on a robust Fos induction. However, under the same treatments, male Kiss1^RP3V^ neurons remained inactive at all times. As expected only the female mice responded with a surge of LH that corresponded to the activation of Kiss1 neurons. One explanation for the lack of a surge in the males might be that the inputs from the SCN are either absent or non-functional.

Nutrition, stress, sensory, and circadian signals are known regulators of reproduction and multiple pathways that directly regulate Kiss1^RP3V^ neurons have been identified, primarily in females. We demonstrate that direct activation of male Kiss1^RP3V^ neurons stimulates the HPG axis, increasing testosterone production, and we speculate that these neurons have a role as physiological regulators of testosterone levels, reproduction, and behavior in male mice. To understand the role of male Kiss1^RP3V^ neurons further studies need to address the afferent pathways that regulate their activation. Studies in female mice demonstrate that the Kiss1^ARC^ neurons send dense projections to the RP3V and directly stimulate burst firing of the Kiss1^RP3V^ neurons by releasing glutamate (Qiu et al., 2016). The stimulated Kiss1^RP3V^ neurons project to the cell bodies of GnRH neurons in the POA and release kisspeptin to activate GnRH neurons; Kiss1^ARC^ neurons project to the terminals of GnRH neurons in the median eminence where they stimulate GnRH release. If these convergent circuits are intact in the male, the Kiss1^RP3V^ neurons could play an essential role in the normal pulsatile regulation of testosterone levels. Nutritional deprivation confers a negative input to Kiss1 neurons and inhibits the HPG axis. Starvation leads to the activation of AgRP neurons in the ARC and these GABAergic AgRP neurons make direct inhibitory synaptic connections with Kiss1 neurons in both the ARC and RP3V (Padilla et al., 2017). This may contribute to the reduced fertility linked to depleted energy stores that can result from either malnutrition or intense athletic training in women. Although much less studied, a similar hypogonadism also occurs in men in response to energy deficit (Wong et al., 2019). The specific role of Kiss1^RP3V^ neurons in this link between nutrition and reproduction is unknown. Our results highlight a potentially novel and unexplored role for Kiss1^RP3V^ neurons in male reproductive physiology and behavior.

## MATERIAL AND METHODS

### Mice

Mice were housed in a temperature and humidity-controlled facility with a 12-h light/dark cycle. All animal care and experimental procedures were approved by the Institutional Animal Care and Use Committee at the University of Washington. *Kiss1*^Cre:GFP^, referred to here as Kiss1-Cre (Gottsch et al., 2011), Ai14 TdTomato Reporter (Madisen et al., 2010) and RiboTag (Sanz et al., 2009) mice were used in this study. All mice used were adults (3 to 4 months old) which had been backcrossed onto a C57BL/6 background. Mice were castrated as described (Gottsch et al., 2011). Briefly, adult male mice were maintained under isoflurane anesthesia while testes were removed through a midline ventral incision. Vasculature to the testes and body wall were sutured, and wound clips were used to close the incision. Mice were given ketoprofen analgesia before surgery and every 24 h after surgery.

### Adeno-associated viral vector delivery

*Kiss1-Cre* male mice were injected bilaterally with a total of 1 μl of AAV1-hSyn-DIO-hM3D(Gq)-mCherry viral vector (7 × 10^11^ viral genomes/ml; 0.5 μl per side) as described previously (Quintana et al., 2012) using the following coordinates: ±0.27 mm mediolateral (ML), +0.145 mm anteroposterior (AP), and −5.5 mm dorsoventral (DV) from Bregma (Paxinos & Franklin, 2013). The vector pAAV-hSyn-DIO-hM3D(Gq)-mCherry was from Bryan Roth (Addgene plasmid #44361; http://n2t.net/addgene:44361; RRID:Addgene_44361) Mice were euthanized 3 to 4 weeks after injections. Kiss1-Cre mice were injected bilaterally with a total of 1 μl of AAV1-Efa1-DIO-Rpl22-3HA-IRES-YFP (AAV-DIO-RiboTag) viral vector (7 × 10^11^ viral genomes/ml; 0.5 μl per side) as previously described (Sanz et al., 2015) using the following coordinates: ±0.27 mm mediolateral (ML), +0.145 mm anteroposterior (AP), and −5.5 mm dorsoventral (DV) from Bregma (Paxinos & Franklin, 2013) Mice were euthanized 3-4 weeks post injection and brain tissue was collected.

### Histology and Imaging

For TdTomato or hM3Dq:mCherry imaging, Kiss1-Cre;Rosa26-flox-stop-TdT or Kiss1-Cre injected with AAV1-DIO-hM3Dq:mCherry mice were euthanized by Beuthanasia-D administration (86 mg/kg pentobarbital sodium, 11 mg/kg phenytoin sodium) followed by cardiac perfusion with 4% paraformaldehyde in PBS, pH 7.4. Brains were dissected and post-fixed at 4°C overnight and cryoprotected in 30% sucrose. Tissue was frozen in dry ice and stored at −80°C until sectioning. Coronal sections (30 μm) containing the RP3V and ARC were used for analysis. Kiss1-Cre;Rosa26-flox-stop-TdT brains were sectioned coronally from +0.5 mm to 0.0 mm Anteroposterior from bregma. Every 4^th^ slice for a total of 4 slices were collected from each of 2 females and 2 males and imaged under a 20X objective on a Zeiss 710 confocal microscope. TdTomato positive cells were counted in an area roughly 736 μm by 736 μm and summed across sections within each animal.

### RiboTag assays

Circular 2 mm-punches containing the RP3V area or the ARC from groups of four RiboTag expressing mice were pooled and homogenized in 1 ml of buffer as described previously (Sanz et al., 2019; Sanz et al., 2015). Briefly, 4 μl of anti-HA antibody (Clone 16B12, BioLegend) were added to 800 μl of the cleared lysate and incubated for 4 h at 4°C. A sample of total lysate (80 μl) was saved as an input. After incubation, 200 μl of protein A/G magnetic beads (Thermo Fisher Scientific) were added and incubated overnight at 4°C with rotation. Immunoprecipitates (IPs) were washed in high salt buffer and RNA from inputs and IPs extracted using the RNeasy micro kit (Qiagen).

### RNA-Seq

#### Library Preparation

NuGEN Ovation RNA-Seq System V2 (Catalog # 7102)(NuGEN, San Carlos, CA) in combination with Ovation Ultralow Library Systems (Catalog #s 0330, 0331) (NuGEN) and 2-10 ng of total RNA were used to make cDNA libraries. Libraries were quantified with NEBNext Library Quant Kit for Illumina (Catalog # E7630, New England BioLabs, Ipswich, MA) before sequencing them on a NextSeq500 (Illumina, San Diego, CA) using NextSeq500 High-Output v2 75 cycles consumables (FC-404-2005; Illumina). Libraries were sequenced 75 cycles using single-ended reads with 6 additional reads for barcodes. Sequencing reads were converted from BCL to FASTQ format and demultiplexed using Bcl2fastq2 (Illumina, 2017). The first 5 base calls were then trimmed and low quality reads filtered out using FASTX-Toolkit (Hannon_Lab, 2014). Reads were aligned to GRCm38/mm10 UCSC mouse genome using TopHat2 (Kim et al., 2013; Trapnell et al., 2009) and quantified and normalized using Cuffnorm from Cufflinks (Trapnell et al., 2010). Data was visualized using R version 3.4.4 “Someone to Lean On” (Gentleman et al., 2018) within RStudio (Rstudio, 2018) utilizing tidyr, dplyr, and ggplot2 (Wickham, 2016; Wickham et al., 2018a; Wickham et al., 2018b; Wickham & Henry, 2018).

### Testosterone assays

Tail vein blood was obtained right before and 1h after clozapine-n-oxide (CNO; 1mg/kg) i.p. administration and allowed to clot in Microtainer serum separator tubes (Beckton-Dickinson) for 1h at RT. Serum was recovered by centrifugation and stored at −80° for later analysis. Testosterone levels were determined using the Testosterone EIA kit (Cayman Chemical, Ann Arbor, MI). Statistical analysis of serum testosterone levels was performed using GraphPad Prism v5.02 software.

## ACKNOWLEDGEMENTS

This work was funded by GM032875 (to GSM) and F32DK116432 (to JCB). MICIU Proyectos I+D+i “Retos Investigacion” (RTI2018-J-100) and Ramòn y Cajal Fellowship (RYC2019-028501-I) to ES. MINECO Proyectos I+D de Excelencia (SAF2017-88108-R) and AGAUR (2017SGR-323) to AQ.

---

**Figure 2–Supplement 1.**
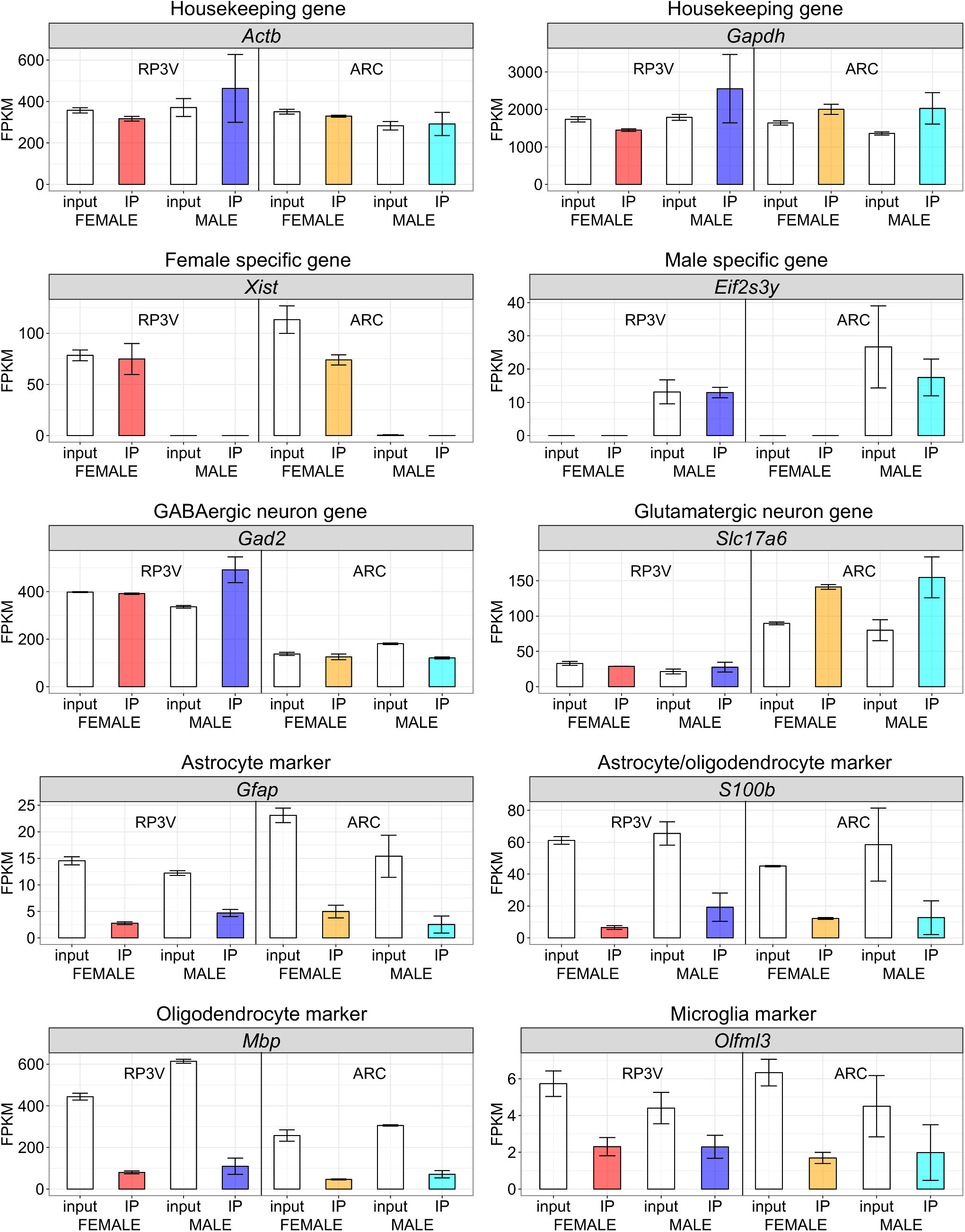
RNA-Seq analysis of ubiquitously expressed “housekeeping” genes, male and female specific genes, GABAergic and Glutamatergic marker genes as well as marker genes for glial cells. The input and IP for RP3V and ARC samples is shown from both male and female mice expressed as normalized FPKM. Two groups of mice (4 animals in each group) were analyzed for each condition and the error bars show the range of the two groups.

**Figure 3–Supplement 1.**
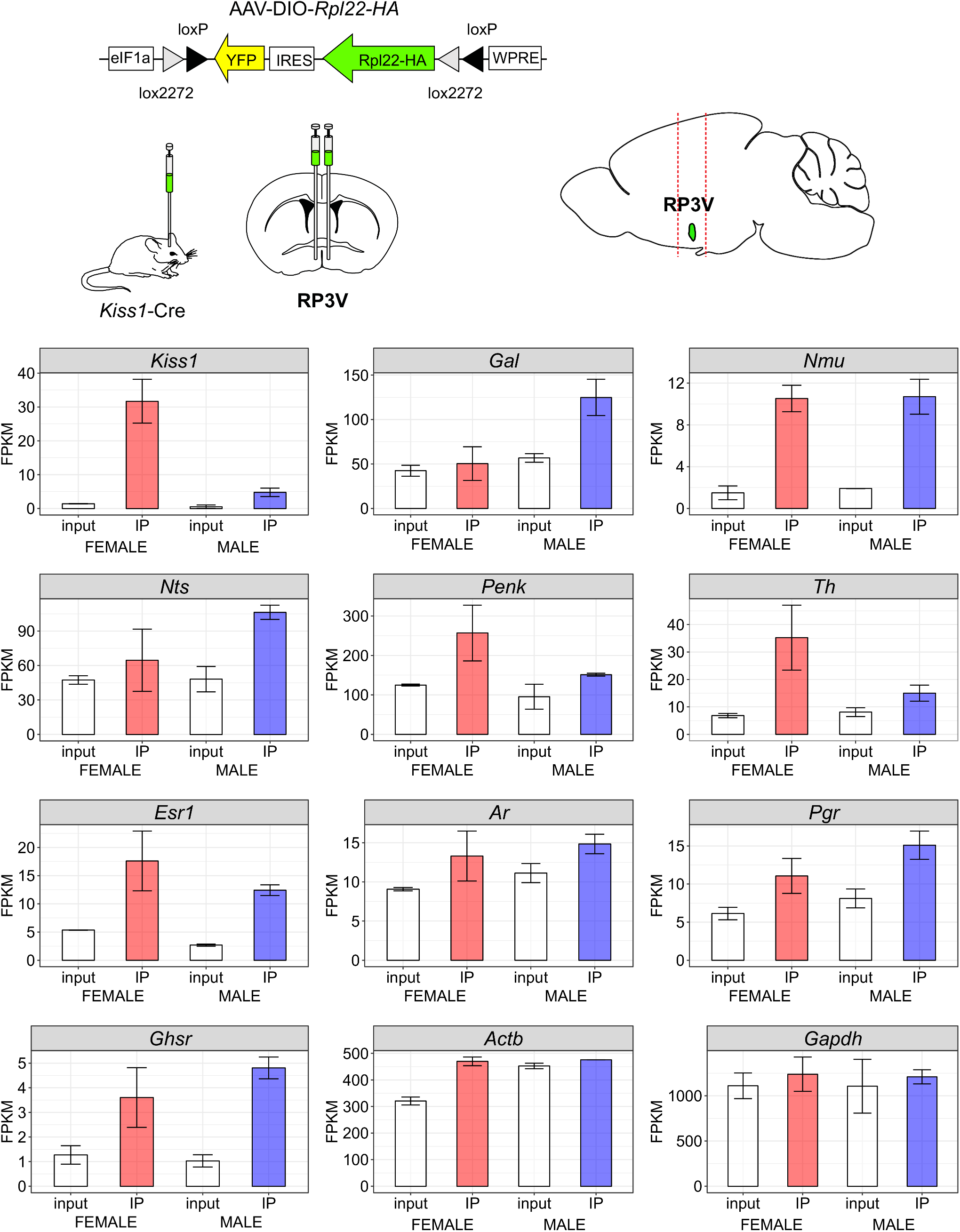
RiboTag analysis after viral injection of AAV-DIO-RiboTag (AAV1-DIO-Rpl22-HA) into Kiss1-Cre mice was used to verify the expression of specific transcripts in adult male Kiss1^RP3V^ neurons. A diagram of the AAV vector is shown that contains a copy of Rpl22-HA (RiboTag) that is activated by Cre recombinase in adult neurons that express Cre recombinase from the Kiss1 locus. Bilateral injection of the viral vector into the RP3V region of Kiss1-Cre mice was followed by a 3 to 4 week recovery before the mice were used to assay gene expression. Two groups of mice (4 animals in each group) were injected with virus and analyzed. Results are shown for the same genes that were analyzed using Kiss1-Cre mice crossed to the RiboTag mouse line shown in Figures 2 and 3. The average values for both Input and IP are shown and the error bar denotes the range of the two groups.

**Figure 3–Supplement 2.**
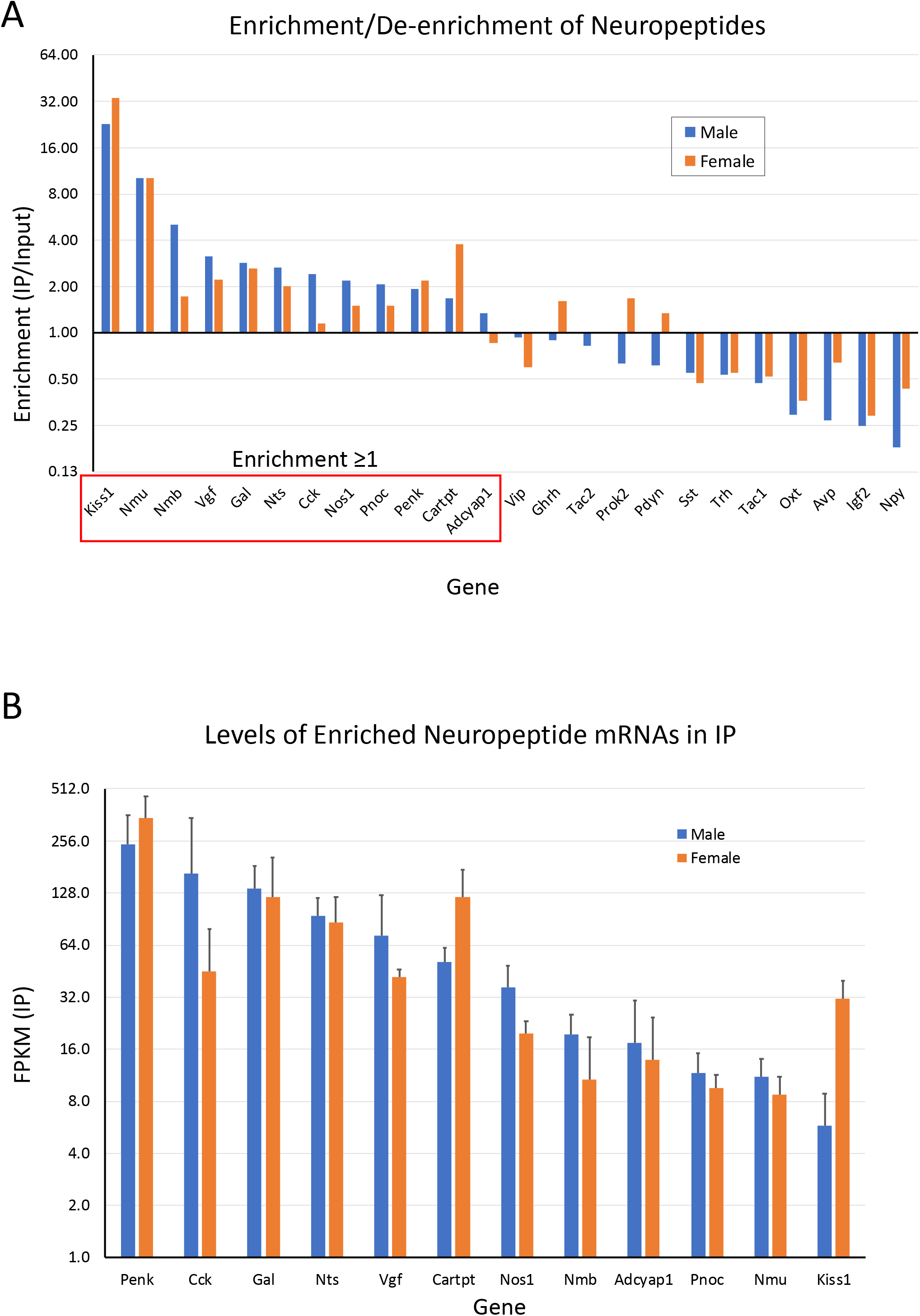
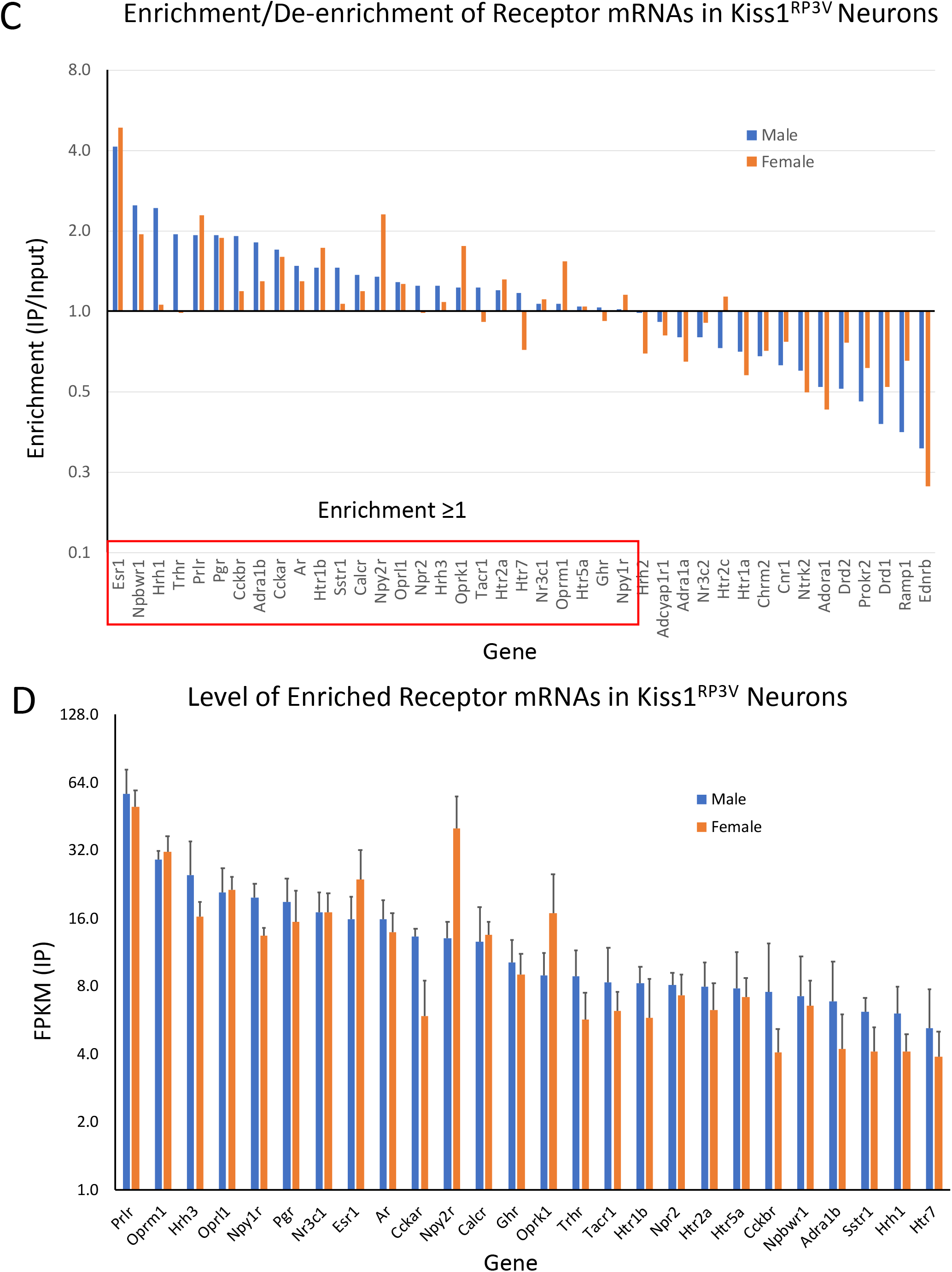
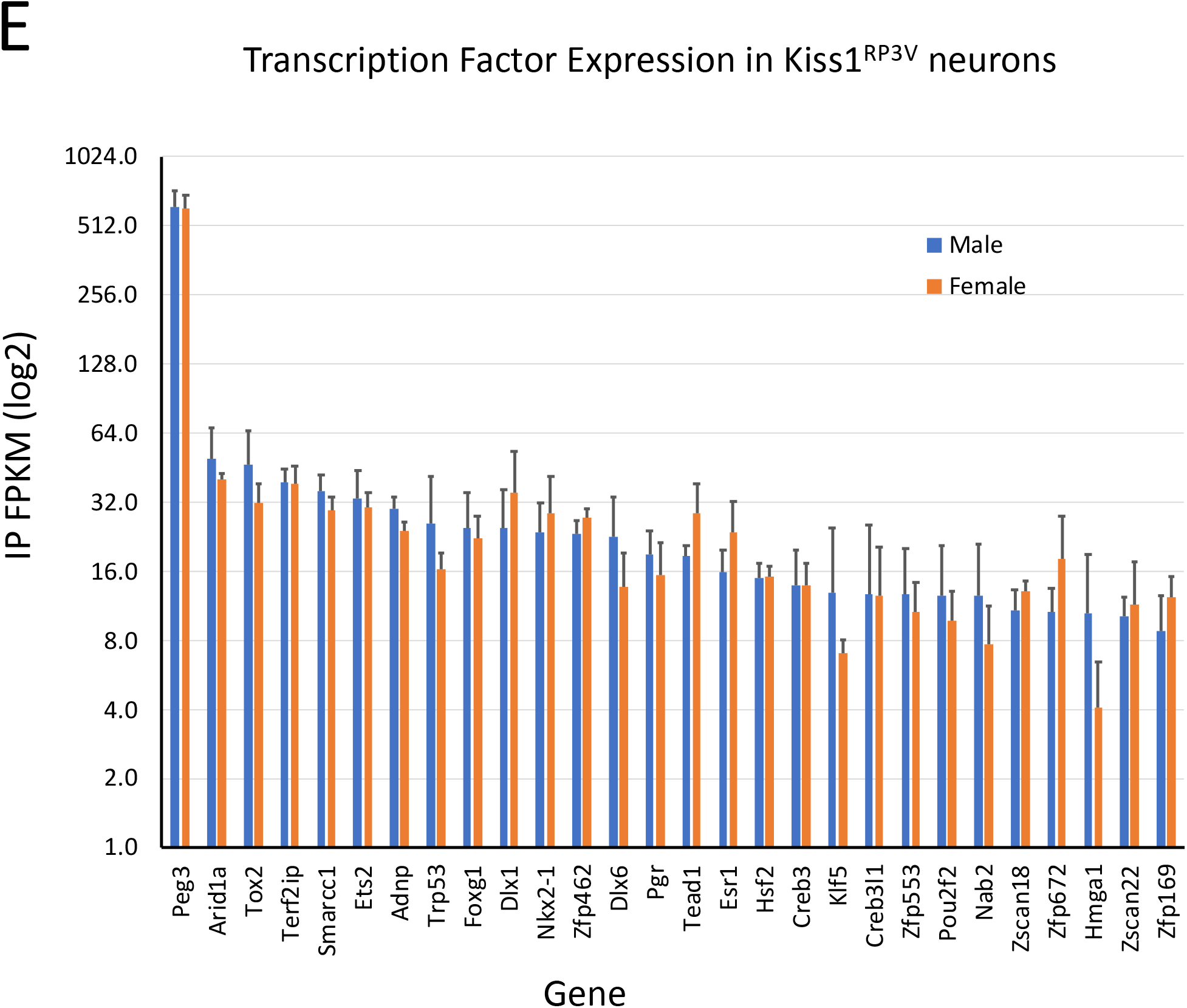
Neuropeptide, receptor, and transcription factor profiles in male and female Kiss1^RP3V^ neurons. A manually curated list of neuropeptide and receptor genes from Moffitt et al (Moffitt et al., 2018) and an atlas of mouse transcription factor genes (Zhou et al., 2017) was used to analyze our RNA-Seq results and derive an unbiased comparison of male and female Kiss1^RP3V^ neuron expression profiles. The RNA-Seq results from the RiboTag mouse and AAV-DIO-RiboTag experiments were normalized together and filtered for mRNAs that showed an averaged expression in either the input (FPKM≥5) or IP (FPKM≥5) of either sex. **(A)** Of the 73 neuropeptides in the Moffitt list, 24 passed the expression filter in the RP3V samples and we then plotted the enrichment (IP/input) for these genes comparing male with female. The enrichment is shown on a log2 scale. **(B)** The 12 neuropeptide mRNAs showing an enrichment ≥1 in males were further analyzed for quantitative expression in the IP samples again comparing male and female Kiss1^RP3V^ neurons. FPKM values are shown on a log2 scale and error bars are the standard deviation (SD) of the 4 samples. **(C)** 139 neuropeptide and hormone receptors were analyzed and 41 passed the filter for expression in either input or IP. The enrichment of these receptors is compared between male and female. **(D)** The 26 receptor mRNAs with enrichment of at least 1 in males were further analyzed to compare the quantitative gene expression in the IP from male and female Kiss1^RP3V^ neurons. The error bars are the SD of the 4 samples. **(E)** Transcription factors that were expressed in either sex at 10 FPKM or greater in the IP and showed an enrichment of 1.5 or greater were compared between male and female Kiss1^RP3V^ neurons. The quantitative results for the 28 transcription factors that passed this filter are shown with the mean and SD.

## REFERENCES

Clarkson, J., & Herbison, A. E. (2006). Postnatal development of kisspeptin neurons in mouse hypothalamus; sexual dimorphism and projections to gonadotropin-releasing hormone neurons. Endocrinology, 147(12), 5817–5825. doi:10.1210/en.2006-0787

Clarkson, J., & Herbison, A. E. (2011). Dual Phenotype Kisspeptin-Dopamine Neurones of the Rostral Periventricular Area of the Third Ventricle Project to Gonadotrophin-Releasing Hormone Neurones. Neuroendocrinology, 23(4), 293–301. doi:10.1111/j.1365-2826.2011.02107.x

Clarkson, J., & Herbison, A. E. (2016). Hypothalamic control of the male neonatal testosterone surge. In Philosophical Transactions of the Royal Society B: Biological Sciences (Vol. 371, pp. 1–9).

Cravo, R. M., Margatho, L. O., Osborne-Lawrence, S., Donato, J., Jr., Atkin, S., Bookout, A. L., Rovinsky, S., Frazao, R., Lee, C. E., Gautron, L., Zigman, J. M., & Elias, C. F. (2011). Characterization of Kiss1 neurons using transgenic mouse models. Neuroscience, 173, 37–56. doi:10.1016/j.neuroscience.2010.11.022

d’Anglemont de Tassigny, X., Fagg, L. A., Dixon, J. P., Day, K., Leitch, H. G., Hendrick, A. G., Zahn, D., Franceschini, I., Caraty, A., Carlton, M. B., Aparicio, S. A., & Colledge, W. H. (2007). Hypogonadotropic hypogonadism in mice lacking a functional Kiss1 gene. Proc Natl Acad Sci U S A, 104(25), 10714–10719. doi:10.1073/pnas.0704114104

de Roux, N., Genin, E., Carel, J. C., Matsuda, F., Chaussain, J. L., & Milgrom, E. (2003). Hypogonadotropic hypogonadism due to loss of function of the KiSS1-derived peptide receptor GPR54. Proc Natl Acad Sci U S A, 100(19), 10972–10976. doi:10.1073/pnas.1834399100

Frazao, R., Dungan Lemko, H. M., da Silva, R. P., Ratra, D. V., Lee, C. E., Williams, K. W., Zigman, J. M., & Elias, C. F. (2014). Estradiol modulates Kiss1 neuronal response to ghrelin. Am J Physiol Endocrinol Metab, 306(6), E606–614. doi:10.1152/ajpendo.00211.2013

Funes, S., Hedrick, J. A., Vassileva, G., Markowitz, L., Abbondanzo, S., Golovko, A., Yang, S., Monsma, F. J., & Gustafson, E. L. (2003). The KiSS-1 receptor GPR54 is essential for the development of the murine reproductive system. Biochem Biophys Res Commun, 312(4), 1357–1363. doi:10.1016/j.bbrc.2003.11.066

Gentleman, R., Ihaka, R., Bates, D., Chambers, J., Dalgaard, P., Hornik, K., Kalibera, T., Lawrence, M., Leisch, F., Ligges, U., Lumley, T., Maechler, M., Morgan, M., Murrell, P., Plummer, M., Ripley, B., Sarkar, D., Lang, D. T., Tierney, L., Urbanek, S., Schwarte, H., Masarotto, G., Iacus, S., Falcon, S., & Murdoch, D. (2018). R (Version 3.5.1) [language and environment for statistical computing]: The R Foundation. Retrieved from https://www.r-project.org/

Gottsch, M. L., Cunningham, M. J., Smith, J. T., Popa, S. M., Acohido, B. V., Crowley, W. F., Seminara, S., Clifton, D. K., & Steiner, R. A. (2004). A role for kisspeptins in the regulation of gonadotropin secretion in the mouse. Endocrinology, 145(9), 4073–4077. doi:10.1210/en.2004-0431

Gottsch, M. L., Popa, S. M., Lawhorn, J. K., Qiu, J., Tonsfeldt, K. J., Bosch, M. A., Kelly, M. J., Rønnekleiv, O. K., Sanz, E., McKnight, G. S., Clifton, D. K., Palmiter, R. D., & Steiner, R. A. (2011). Molecular Properties of Kiss1 Neurons in the Arcuate Nucleus of the Mouse. Endocrinology, 152, 4298–4309. doi:10.1210/en.2011-1521

Hannon_Lab, G. J. (2014). FASTX-Toolkit (Version 0.0.14): Hannon Lab. Retrieved from http://hannonlab.cshl.edu/fastx_toolkit/download.html

Homma, T., Sakakibara, M., Yamada, S., Kinoshita, M., Iwata, K., Tomikawa, J., Kanazawa, T., Matsui, H., Takatsu, Y., Ohtaki, T., Matsumoto, H., Uenoyama, Y., Maeda, K., & Tsukamura, H. (2009). Significance of neonatal testicular sex steroids to defeminize anteroventral periventricular kisspeptin neurons and the GnRH/LH surge system in male rats. Biol Reprod, 81(6), 1216–1225. doi:10.1095/biolreprod.109.078311

Illumina. (2017). bcl2fastq2 (Version (Version 2.2)): Illumina. Retrieved from https://support.illumina.com/downloads/bcl2fastq-conversion-software-v2-20.html.

Kauffman, A. S. (2010). Gonadal and nongonadal regulation of sex differences in hypothalamic Kiss1 neurones. J Neuroendocrinol, 22(7), 682–691. doi:10.1111/j.1365-2826.2010.02030.x

Kim, D., Pertea, G., Trapnell, C., Pimentel, H., Kelley, R., & Salzberg, S. L. (2013). TopHat2: accurate alignment of transcriptomes in the presence of insertions, deletions and gene fusions. Genome biology, 14(4), R36. doi:10.1186/gb-2013-14-4-r36

Krashes, M. J., Koda, S., Ye, C., Rogan, S. C., Adams, A. C., Cusher, D. S., Maratos-Flier, E., Roth, B. L., & Lowell, B. B. (2011). Rapid, reversible activation of AgRP neurons drives feeding behavior in mice. J Clin Invest, 121(4), 1424–1428. doi:10.1172/JCI46229

Kumar, D., Candlish, M., Periasamy, V., Avcu, N., Mayer, C., & Boehm, U. (2015). Specialized subpopulations of kisspeptin neurons communicate with GnRH neurons in female mice. Endocrinology, 156(1), 32–38. doi:10.1210/en.2014-1671

Madisen, L., Zwingman, T. A., Sunkin, S. M., Oh, S. W., Zariwala, H. A., Gu, H., Ng, L. L., Palmiter, R. D., Hawrylycz, M. J., Jones, A. R., Lein, E. S., & Zeng, H. (2010). A robust and high-throughput Cre reporting and characterization system for the whole mouse brain. Nat Neurosci, 13(1), 133–140. doi:10.1038/nn.2467

Moffitt, J. R., Bambah-Mukku, D., Eichhorn, S. W., Vaughn, E., Shekhar, K., Perez, J. D., Rubinstein, N. D., Hao, J., Regev, A., Dulac, C., & Zhuang, X. (2018). Molecular, spatial, and functional single-cell profiling of the hypothalamic preoptic region. Science, 362(6416), eaau5324. doi:10.1126/science.aau5324

Navarro, V. M., Gottsch, M. L., Wu, M., Garcia-Galiano, D., Hobbs, S. J., Bosch, M. A., Pinilla, L., Clifton, D. K., Dearth, A., Ronnekleiv, O. K., Braun, R. E., Palmiter, R. D., Tena-Sempere, M., Alreja, M., & Steiner, R. A. (2011). Regulation of NKB pathways and their roles in the control of Kiss1 neurons in the arcuate nucleus of the male mouse. Endocrinology, 152(11), 4265–4275. doi:10.1210/en.2011-1143

Nestor, C. C., Qiu, J., Padilla, S. L., Zhang, C., Bosch, M. A., Fan, W., Aicher, S. A., Palmiter, R. D., Ronnekleiv, O. K., & Kelly, M. J. (2016). Optogenetic Stimulation of Arcuate Nucleus Kiss1 Neurons Reveals a Steroid-Dependent Glutamatergic Input to POMC and AgRP Neurons in Male Mice. Mol Endocrinol, 30(6), 630–644. doi:10.1210/me.2016-1026

Oakley, A. E., Clifton, D. K., & Steiner, R. A. (2009). Kisspeptin signaling in the brain. Endocr Rev, 30(6), 713–743. doi:10.1210/er.2009-0005

Padilla, S. L., Qiu, J., Nestor, C. C., Zhang, C., Smith, A. W., Whiddon, B. B., Ronnekleiv, O. K., Kelly, M. J., & Palmiter, R. D. (2017). AgRP to Kiss1 neuron signaling links nutritional state and fertility. Proc Natl Acad Sci U S A, 114(9), 2413–2418. doi:10.1073/pnas.1621065114

Paxinos, G., & Franklin, K. B. J. (2013). The Mouse Brain in Stereotaxic Coordinates. San Diego, CA: Academic Press.

Piet, R., Kalil, B., McLennan, T., Porteous, R., Czieselsky, K., & Herbison, A. E. (2018). Dominant Neuropeptide Cotransmission in Kisspeptin-GABA Regulation of GnRH Neuron Firing Driving Ovulation. J Neurosci, 38(28), 6310–6322. doi:10.1523/JNEUROSCI.0658-18.2018

Poling, M. C., Luo, E. Y., & Kauffman, A. S. (2017). Sex Differences in Steroid Receptor Coexpression and Circadian-Timed Activation of Kisspeptin and RFRP-3 Neurons May Contribute to the Sexually Dimorphic Basis of the LH Surge. Endocrinology, 158(10), 3565–3578. doi:10.1210/en.2017-00405

Porteous, R., Petersen, S. L., Yeo, S. H., Bhattarai, J. P., Ciofi, P., de Tassigny, X. D., Colledge, W. H., Caraty, A., & Herbison, A. E. (2011). Kisspeptin neurons co-express met-enkephalin and galanin in the rostral periventricular region of the female mouse hypothalamus. J Comp Neurol, 519(17), 3456–3469. doi:10.1002/cne.22716

Qiu, J., Nestor, C. C., Zhang, C., Padilla, S. L., Palmiter, R. D., Kelly, M. J., & Ronnekleiv, O. K. (2016). High-frequency stimulation-induced peptide release synchronizes arcuate kisspeptin neurons and excites GnRH neurons. eLife, 5, 3–5. doi:10.7554/eLife.16246

Qiu, J., Rivera, H. M., Bosch, M. A., Padilla, S. L., Stincic, T. L., Palmiter, R. D., Kelly, M. J., & Ronnekleiv, O. K. (2018). Estrogenic-dependent glutamatergic neurotransmission from kisspeptin neurons governs feeding circuits in females. eLife, 7, 1–60. doi:10.7554/eLife.35656

Quintana, A., Sanz, E., Wang, W., Storey, G. P., Guler, A. D., Wanat, M. J., Roller, B. A., La Torre, A., Amieux, P. S., McKnight, G. S., Bamford, N. S., & Palmiter, R. D. (2012). Lack of GPR88 enhances medium spiny neuron activity and alters motor- and cue-dependent behaviors. Nat Neurosci, 15(11), 1547–1555. doi:10.1038/nn.3239

RStudio. (2018). RStudio (Version 1.1.463) [integrated development environment (IDE)]: RStudio. Retrieved from https://www.rstudio.com/products/rstudio/download/#download

Sanz, E., Bean, J. C., Carey, D. P., Quintana, A., & McKnight, G. S. (2019). RiboTag: Ribosomal Tagging Strategy to Analyze Cell-Type-Specific mRNA Expression In Vivo. Curr Protoc Neurosci, 88(1), e77. doi:10.1002/cpns.77

Sanz, E., Quintana, A., Deem, J. D., Steiner, R. A., Palmiter, R. D., & McKnight, G. S. (2015). Fertility-regulating Kiss1 neurons arise from hypothalamic POMC-expressing progenitors. J Neurosci, 35(14), 5549–5556. doi:10.1523/JNEUROSCI.3614-14.2015

Sanz, E., Yang, L., Su, T., Morris, D. R., McKnight, G. S., & Amieux, P. S. (2009). Cell-type-specific isolation of ribosome-associated mRNA from complex tissues. Proc Natl Acad Sci U S A, 106(33), 13939–13944. doi:10.1073/pnas.0907143106

Semaan, S. J., & Kauffman, A. S. (2013). Emerging concepts on the epigenetic and transcriptional regulation of the Kiss1 gene. Int J Dev Neurosci, 31(6), 452–462. doi:10.1016/j.ijdevneu.2013.03.006

Seminara, S. B., Messager, S., Chatzidaki, E. E., Thresher, R. R., Acierno, J. S., Jr., Shagoury, J. K., Bo-Abbas, Y., Kuohung, W., Schwinof, K. M., Hendrick, A. G., Zahn, D., Dixon, J., Kaiser, U. B., Slaugenhaupt, S. A., Gusella, J. F., O’Rahilly, S., Carlton, M. B., Crowley, W. F., Jr., Aparicio, S. A., & Colledge, W. H. (2003). The GPR54 gene as a regulator of puberty. N Engl J Med, 349(17), 1614–1627. doi:10.1056/NEJMoa035322

Simerly, R. B., Zee, M. C., Pendleton, J. W., Lubahn, D. B., & Korach, K. S. (1997). Estrogen receptor-dependent sexual differentiation of dopaminergic neurons in the preoptic region of the mouse. Proc Natl Acad Sci U S A, 94(25), 14077–14082. doi:10.1073/pnas.94.25.14077

Smith, J. T., Dungan, H. M., Stoll, E. A., Gottsch, M. L., Braun, R. E., Eacker, S. M., Clifton, D. K., & Steiner, R. A. (2005). Differential regulation of KiSS-1 mRNA expression by sex steroids in the brain of the male mouse. Endocrinology, 146(7), 2976–2984. doi:10.1210/en.2005-0323

Stephens, S. B., Tolson, K. P., Rouse, M. L., Jr., Poling, M. C., Hashimoto-Partyka, M. K., Mellon, P. L., & Kauffman, A. S. (2015). Absent Progesterone Signaling in Kisspeptin Neurons Disrupts the LH Surge and Impairs Fertility in Female Mice. Endocrinology, 156(9), 3091–3097. doi:10.1210/en.2015-1300

Trapnell, C., Pachter, L., & Salzberg, S. L. (2009). TopHat: discovering splice junctions with RNA-Seq. Bioinformatics, 25(9), 1105–1111. doi:10.1093/bioinformatics/btp120

Trapnell, C., Williams, B. A., Pertea, G., Mortazavi, A., Kwan, G., van Baren, M. J., Salzberg, S. L., Wold, B. J., & Pachter, L. (2010). Transcript assembly and quantification by RNA-Seq reveals unannotated transcripts and isoform switching during cell differentiation. Nature biotechnology, 28(5), 511–515. doi:10.1038/nbt.1621

Wang, L., & Moenter, S. M. (2020). Differential Roles of Hypothalamic AVPV and Arcuate Kisspeptin Neurons in Estradiol Feedback Regulation of Female Reproduction. Neuroendocrinology, 110(3-4), 172–184. doi:10.1159/000503006

Wang, L., Vanacker, C., Burger, L. L., Barnes, T., Shah, Y. M., Myers, M. G., & Moenter, S. M. (2019). Genetic dissection of the different roles of hypothalamic kisspeptin neurons in regulating female reproduction. eLife, 8, 1–26. doi:10.7554/eLife.43999

Wickham, H. (2016). ggplot2: *Elegant Graphics for Data Analysis*. New York, NY: Springer-Verlag New York.

Wickham, H., Chang, W., Henry, L., Pedersen, T. L., Takahashi, K., Wilke, C., Woo, K., & Yutani, H. (2018a). ggplot2 (Version 3.1.0). Retrieved from https://ggplot2.tidyverse.org/

Wickham, H., François, R., Henry, L., & Müller, K. (2018b). dplyr (Version 0.7.8): RStudio. Retrieved from https://dplyr.tidyverse.org/

Wickham, H., & Henry, L. (2018). tidyr (Version 0.8.2.9000): RStudio. Retrieved from https://tidyr.tidyverse.org/

Williams, W. P., 3rd, Jarjisian, S. G., Mikkelsen, J. D., & Kriegsfeld, L. J. (2011). Circadian control of kisspeptin and a gated GnRH response mediate the preovulatory luteinizing hormone surge. Endocrinology, 152(2), 595–606. doi:10.1210/en.2010-0943

Wong, H. K., Hoermann, R., & Grossmann, M. (2019). Reversible male hypogonadotropic hypogonadism due to energy deficit. Clin Endocrinol (Oxf), 91(1), 3–9. doi:10.1111/cen.13973

Zhang, C., Bosch, M. A., Qiu, J., Ronnekleiv, O. K., & Kelly, M. J. (2015). 17beta-Estradiol increases persistent Na(+) current and excitability of AVPV/PeN Kiss1 neurons in female mice. Mol Endocrinol, 29(4), 518–527. doi:10.1210/me.2014-1392

Zhang, C., Tonsfeldt, K. J., Qiu, J., Bosch, M. A., Kobayashi, K., Steiner, R. A., Kelly, M. J., & Ronnekleiv, O. K. (2013). Molecular mechanisms that drive estradiol-dependent burst firing of Kiss1 neurons in the rostral periventricular preoptic area. Am J Physiol Endocrinol Metab, 305(11), E1384–1397. doi:10.1152/ajpendo.00406.2013

Zhou, Q., Liu, M., Xia, X., Gong, T., Feng, J., Liu, W., Liu, Y., Zhen, B., Wang, Y., Ding, C., & Qin, J. (2017). A mouse tissue transcription factor atlas. Nat Commun, 8, 15089. doi:10.1038/ncomms15089

